# α-proteobacterial RNA degradosomes assemble liquid-liquid phase separated RNP bodies

**DOI:** 10.1101/272286

**Authors:** Nadra Al-Husini, Dylan T. Tomares, W. Seth Childers, Jared M. Schrader

## Abstract

Ribonucleoprotein (RNP) granules play an important role in organizing eukaryotic mRNA metabolism via liquid-liquid phase separation (LLPS) of mRNA decay factors into membrane-less “droplet” organelles in the cytoplasm. Here we show that the bacterium *Caulobacter crescentus* Ribonuclease (RNase) E assembles RNP LLPS droplets that we term <u>b</u>acterial <u>R</u>NP-bodies (BR-bodies) similar to eukaryotic P-bodies and stress granules. RNase E requires RNA to assemble a BR-body, and disassembly requires RNA cleavage, suggesting BR-bodies provide localized sites of RNA degradation. The unstructured C-terminal domain of RNase E is both necessary and sufficient to assemble the core of the BR-body, is functionally conserved in related α-proteobacteria, and influences mRNA degradation. BR-bodies are rapidly induced under cellular stresses and provide enhanced cell growth under stress. To our knowledge, *Caulobacter* RNase E is the first bacterial protein identified that forms LLPS droplets, providing an effective strategy for subcellular organization in cells lacking membrane bound compartments.

## Introduction

mRNA turnover provides an essential pathway in the control of gene expression across the domains of life. The most common bacterial protein controlling mRNA turnover is RNase E which is conserved in approximately half of all bacteria (Ait-Bara and Carpousis, 2015; Mohanty and Kushner, 2016). In the γ-proteobacteria *E. coli* (*Eco* henceforth) RNase E is a membrane tethered protein whose membrane anchoring sequence is required for its ability to localize into short-lived membrane anchored foci (Khemici et al., 2008; Strahl et al., 2015; Taghbalout and Rothfield, 2007). In the α-proteobacteria *Caulobacter crescentus* (*Ccr* henceforth) RNase E was instead found to localize into patchy foci within the nucleoid filled cytoplasm (Bayas et al., 2018; Montero Llopis et al., 2010). The *Ccr* RNase E protein also scaffolds an RNA degradosome multi-protein complex containing the DEAD-box RNA helicase RhlB, the metabolic enzyme Aconitase, the exonuclease PNPase, and the exonuclease RNase D (Hardwick et al., 2011; Voss et al., 2014). The functional role of *Ccr*’s cytoplasmic RNase E foci and their role in the subcellular organization of the *Ccr* RNA degradosome has remained unclear.

In eukaryotes, enzymes involved in mRNA turnover are organized in discrete cytoplasmic ribonucleoprotein (RNP) granules termed processing-bodies (P-bodies) or stress granules where mRNAs can be stored or degraded (Parker and Sheth, 2007). Interestingly, P-bodies and stress granules contain Xrn1, an eukaryotic exonuclease whose catalytic activity controls mRNA turnover (Kedersha et al., 2005), a key cellular function that is shared by RNase E (Hammarlof et al., 2015; Ono and Kuwano, 1979). P-bodies are conserved in eukaryotes from yeast to humans and require intrinsically disordered domains to facilitate assembly into discrete cytoplasmic liquid-liquid phase separated (LLPS) droplets (Courchaine et al., 2016; Parker and Sheth, 2007). Importantly, LLPS RNP bodies provide membrane-less intracellular compartments to organize the mRNA turnover process in the eukaryotic cytoplasm and have been referred to as “droplet organelles” (Courchaine et al., 2016). Such droplet organelles are thought to exist in other cellular contexts such as nucleoli, cajal bodies, stress granules, and nuclear bodies, yet no such structures have been identified in bacterial cells (Brangwynne, 2013; Courchaine et al., 2016).

Here we provide evidence that *Ccr* RNase E assembles <u>b</u>acterial <u>R</u>NP bodies (BR-bodies) which share many similarities with eukaryotic P-bodies and stress granules. *Ccr* RNase E forms RNA dependent bodies via its intrinsically disordered CTD which is both necessary and sufficient for BR-body assembly. Alternating blocks of positive and negative charges are needed for self-assembly and present across all α-proteobacterial CTDs where BR-bodies are functionally conserved. *Ccr* RNase E BR-bodies are dynamic under log-growth conditions and RNA cleavage by RNase E is required for their dissociation, suggesting BR-bodies provide active sites of RNA cleavage. Similar to P-bodies or stress granules *Ccr* RNase E can also assemble into LLPS droplets that recruit other mRNA decay factors into BR-bodies. BR-bodies likely play an important role in facilitating mRNA decay as BR-bodies compete with translating ribosomes for mRNA substrates and RNase E mutants lacking the CTD are defective in degradation of an RNase E controlled mRNA. Finally, the assembly of RNase E based BR-bodies are rapidly induced by certain cell stresses and the ability to assemble BR-bodies gives *Ccr* cells enhanced stress resistance. The identification of BR-bodies suggests that the organization of mRNA degradation machinery into membrane-less droplet organelles is more widespread than previously appreciated. To our knowledge, this is the first bacterial body found to be assembled by LLPS, and LLPS may provide an important general mechanism for organizing biochemical reactions in bacteria as they lack membrane bound organelles.

## Results

### mRNA dependent foci formation of α-proteobacterial RNase E

To investigate whether *Ccr* RNase E shares the ability to assemble RNP granules like the eukaryotic Xrn1 a RNase E-YFP fusion strain was visualized under log growth conditions and after depletion of cellular mRNA. In line with a previous report (Montero Llopis et al., 2010), *Ccr* RNase E-YFP localizes into patchy foci within the cytoplasm (Fig 1A). Across a population of cells, cells of all stages of the cell cycle contain these patchy foci with significant cell to cell variability in position, number, and intensity of foci (Fig 1A). The observed foci do not appear to be a result of the YFP tag’s propensity to aggregate (Landgraf et al., 2012), as monomeric superfolder GFP tagged RNase E localizes into similar patchy foci (Fig S1). To examine whether the presence of mRNA was required for the RNase E foci to form, cells were treated with high levels of rifampicin (100μg/mL) to rapidly deplete cellular mRNAs without disrupting the protein level of RNase E-YFP. Treatment of rifampicin for 30 minutes significantly diminished RNase E-YFP fluorescent foci while the total cell intensity of YFP signal remains similar (Fig 1A, Fig S2) (Bayas et al., 2018). The loss in RNase E-YFP foci upon rifampicin treatment suggests that mRNA is indeed required for RNase E foci to assemble.

**Figure 1.**
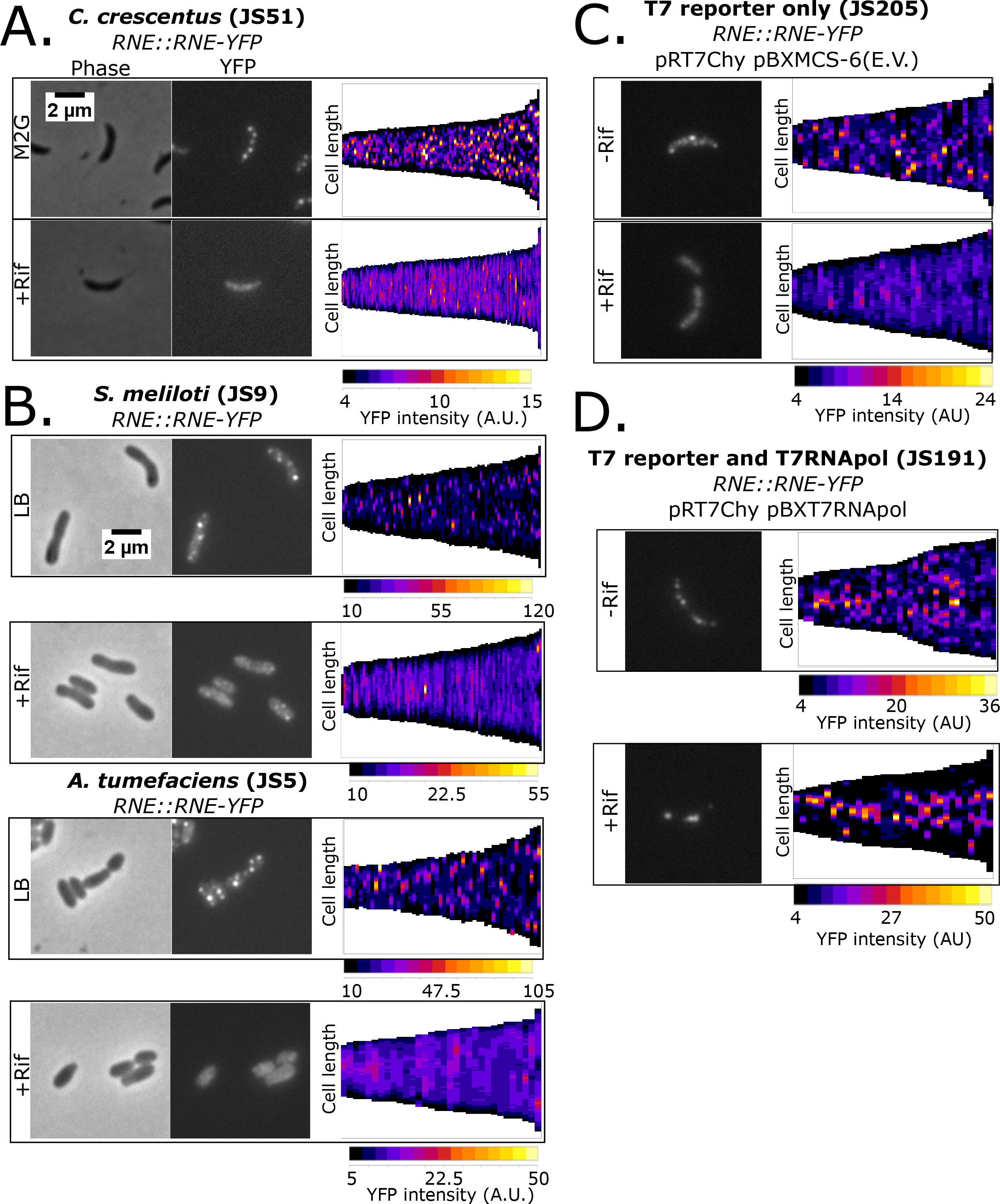
α-proteobacterial RNase E assembles mRNA dependent foci. A.) Localization patterns for NA1000 RNase E – YFP strain(JS51) grown in M2G minimal media and cells were imaged on an M2G 1.5% agarose pad. Demograph displays the YFP intensity as a heat map across the long axis of single cells as a vertical slice. The cells were arranged per their lengths (demograph plot generated with microbeJ). Untreated cells n=84. Rifampicin (100 μg/mL) pre-treated cells n= 149. B.) Localization patterns for log phase *Sme 1021* or *Atu C58* RNase E – YFP strains (JS9 and JS5, respectively). Cells were pre-treated with rifampicin (100 μg/mL) for 30 minutes prior to spotting on the pad as indicated. n=78 *Sme* untreated, n=83 *Sme* rifampicin treated. n=52 *Atu* untreated, and n=32 *Atu* rifampicin treated. C.) Localization patterns for *Ccr* cells expressing RNase E – YFP containing a T7 polymerase driven mCherry reporter plasmid (pRT7Chy) and a pBX empty vector plasmid (pBXMCS-6) (JS205). Cells were grown in log phase in PYE medium with 0.2% xylose and imaged on an M2G 1.5% agarose pad. Cells were either not treated or pre-treated with rifampicin (100 μg/mL) for 30 minutes prior to spotting on the pad and imaging. To the right is a demograph of YFP intensity profile along the long axis of many cells (generated with microbeJ) n=39 untreated, n=39 rif-treated. D.) Localization patterns for *Ccr* cells expressing RNase E – YFP containing a T7 polymerase driven mCherry reporter plasmid (pRT7Chy) and a pBX plasmid containing the rif^R^ T7 RNA polymerase (pBXT7RNApol) (JS191). Cells were grown identical to panel C. To the right is a demograph of YFP intensity profile along the long axis of many cells (generated with microbeJ) n=38 untreated, and n=32 rifampicin treated.

RNase E is conserved in all α-proteobacterial genomes and its ability to assemble BR-bodies in other α-proteobacteria was examined. RNase E-YFP fusions in *Sinorhizobium meliloti(Sme* henceforth) and *Agrobacterium tumefaciens (Atu* henceforth) were generated and their subcellular localization patterns examined. *Sme* and *Atu* RNase E-YFP yielded a similar pattern of YFP foci to that of *Ccr* (Fig 1A,B). Of note, there appear to be fewer RNase E-YFP foci per cell in *Sme* than in *Ccr* or *Atu* (Fig 1B, Fig S3). The foci range in size from a median width of 0.2μm for *Ccr* to 0.4 μm in *Sme* and *Atu*, all on the same size scale as eukaryotic P-bodies or stress granules (Wheeler et al., 2016). Importantly, both *Atu* and *Sme* RNase E foci disperse after a 30-minute treatment with high levels of rifampicin (100 μg/mL), suggesting RNase E-YFP foci also require mRNA in *Atu* and *Sme* (Fig 1).

In addition to its blockage of transcription and subsequent depletion of cellular mRNA, rifampicin treatment also disrupts the chromosome structure (Le et al., 2013). Therefore, rifampicin treatment data alone does not distinguish which cellular effect (mRNA depletion or disruption of chromosome structure) was responsible for the abolishment of the RNase E YFP foci. To determine which cellular effect is the cause of the loss of RNase E foci, the rifampicin resistant T7 RNA polymerase system was utilized to produce mRNAs in the presence of rifampicin (Strahl et al., 2015). The rifampicin resistant T7 RNA polymerase was introduced into *Ccr* on a high copy pBX replicating plasmid together with a low copy replicating plasmid containing a T7 driven mCherry gene and a chromosomally tagged RNase E-YFP (Fig S4). *Ccr* cells harboring the T7 promoter mCherry plasmid and a pBX-empty vector plasmid had rifampicin sensitive RNase E foci as observed in the wild type strain (Fig 1C, Fig S5). In cells containing both the T7 promoter driven mCherry plasmid and the pBX T7 RNA polymerase containing plasmids, robust RNase E foci with increased numbers were observed (Fig 1D, Fig S5), likely due to higher mRNA concentration in the cells. Upon treatment with rifampicin, which disrupts the chromosome structure but does not affect transcription of the T7 driven mCherry mRNA, robust RNase E foci were observed composed of nearly all the RNase E in the cell with localization near the 1/3 and 2/3 points along the long cell axis (Fig 1D, Fig S5) where the reporter plasmids have been found to localize (Kahng and Shapiro, 2003). These data show that the abolishment of RNase E foci by rifampicin is a result of the depletion of cellular mRNAs and not the disruption of the chromosome structure.

### The intrinsically disordered CTD is necessary and sufficient for LLPS

*Ccr* RNase E is composed of two major protein domains, the catalytic N-terminal Domain (NTD) which is essential for cellular viability (Christen et al., 2011) and its sequence is highly conserved across bacterial species, and a poorly conserved CTD (Fig 2A, Fig S6). The NTD contains the S1 RNA binding domain which unwinds folded mRNA substrates near the active site, the active site within the DNase I domain and the 5’ monophosphate sensor domain, while the CTD contains an intrinsically disordered region with conserved Arg-Rich RNA binding sites (Fig S6). When the CTD region was deleted from RNase E the protein became diffuse throughout the cytoplasm (Fig 2A, Fig S7,S8), showing that the CTD is necessary for assembly into foci. In contrast, when the NTD was deleted robust foci formation was observed as in the full-length protein (Fig 2A, Fig S7,S8).

**Figure 2.**
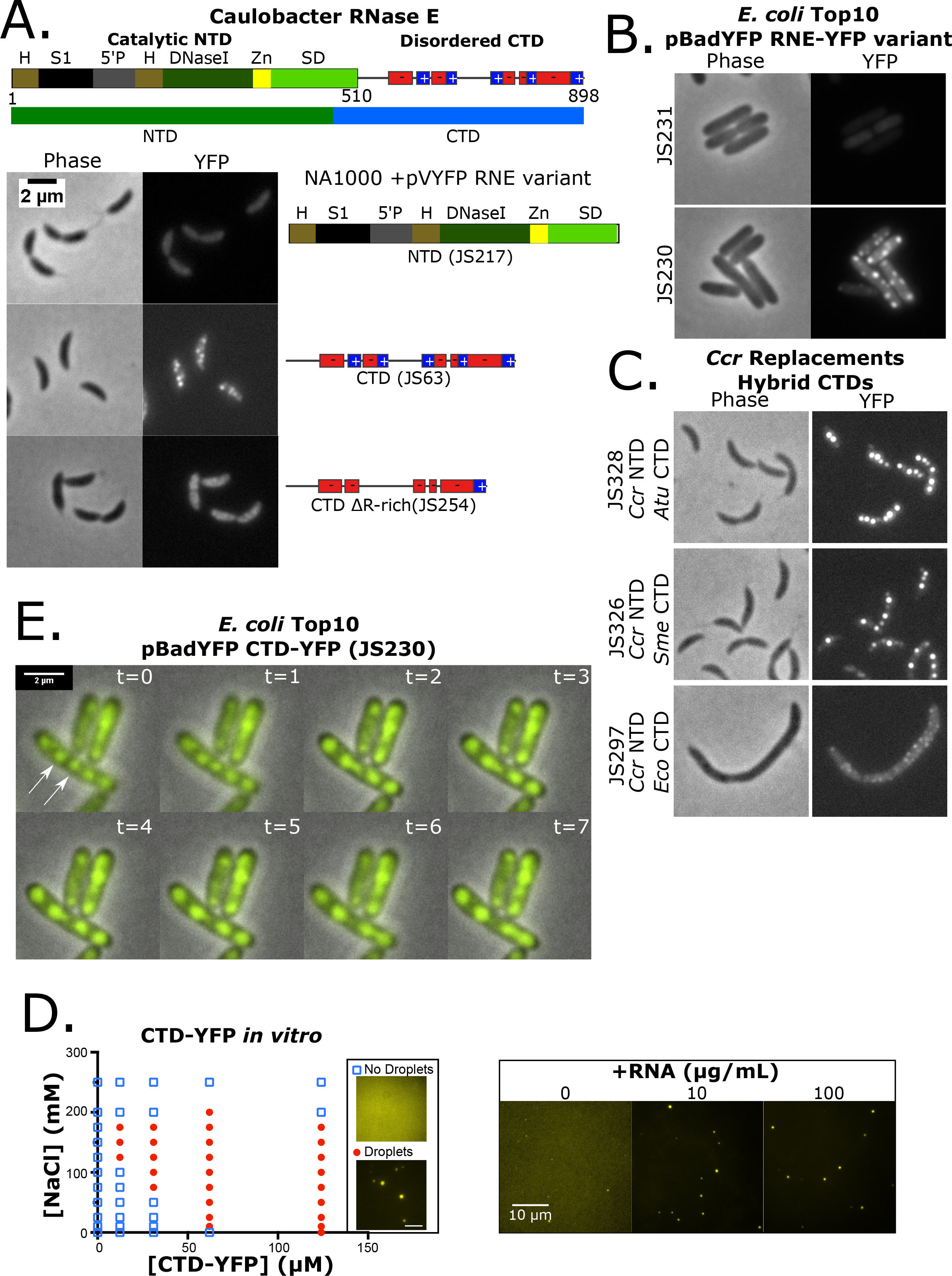
α-proteobacterial RNase E CTD is required for BR-body assembly into liquid-liquid phase separated droplets. A.) Domain architecture for the *Ccr* RNase E protein (Hardwick et al., 2011). The catalytic N-terminal domain (NTD, 1-510 amino acids) and disordered C-terminal domain (CTD, 451-898) are highlighted. H= RNase H domain, S1= ribosomal protein S1 domain, 5’P = 5’ monophosphate sensor domain, DNaseI= Catalytic region, Zn = Zinc link, SD= small domain, and AR= Arginine rich RNA binding sites. Based on the conserved Pfam (PF10150) RNase EG domain (residues 178-450) RNase E was split into the NTD (residues 1-450) and the CTD (residues 451-898). Positive and negative charged patches are indicated with red and blue boxes. Subcellular localization patterns in *Ccr* of the YFP tagged NTD(JS217), CTD (JS63), and CTD ΔR-rich (JS254) mutants of RNase E as indicated. Log phase cells were grown in PYE/gentamycin, induced with 0.5mM vanillate for 6 hours and imaged on M2G 1.5% agarose pads. B.) Heterologous subcellular localization patterns of *Ccr* RNase E in *Eco* top10 cells with the YFP tagged mutants of RNase E as indicated (JS230 and JS231). Log phase cells were grown in LB/ampicillin at 37°C, induced with 0.0004% arabinose for 1 hour and imaged on M2G 1.5% agarose pads. C.) RNase E CTD domain fusions with the CTD domains from *Atu*, *Sme*, or *Eco* fused to the *Ccr* NTD (Domain architectures based on interpro). Subcellular localization of RNase E-YFP CTD fusions were then monitored in RNase E replacement strains. All strains were grown under replacement conditions and imaged in log phase on M2G 1.5% agarose pads. D.) Phase diagram of purified *Ccr* RNase E CTD-YFP (purified protein was made from JS244 cells) incubated at the indicated concentrations of protein and NaCl. LLPS droplets phase boundary is indicated by red circles. Right, droplets were formed with addition of the indicated concentrations of poly-A RNA at 100mM NaCl and 12.4 μM CTD-YFP. E.) Time-lapse images of JS230 showing droplet fusion. Images were taken at 1 minute intervals and the white arrows indicate the two BR-bodies that fuse.

One striking feature of the disordered C-terminus is that the charged residues are organized as clustered blocks of negative and positive charge across the CTD (Fig 2A, Fig S9). Each block ranges in length of 10-30 residues long with a sliding net charge per 10 amino acids that range +2 to +8 for positively charged blocks and −2 to −8 for negatively charged blocks. Such amino acid organization within an intrinsically disordered protein is similar to the primary constituent of eukaryotic germ granules Dd×4 (Nott et al., 2015). The disruption of Ddx4’s charge patterning resulted in a loss of LLPS, suggesting that the multivalent electrostatic protein-protein interactions play an important role in self-assembly (Nott et al., 2015). To test the importance of charged blocks, the positively charged blocks were deleted from the CTD (ΔR-rich) which resulted in diffuse localization of CTD-YFP. This suggests that charged blocks and/or RNA binding capabilities are critical for foci formation (Fig 2A). *Eco* cells are evolutionarily distinct from *Ccr* and provide an *in vivo* test tube for the examination of the intrinsic protein localization pattern of proteins. In line with the results in *Ccr* cells, expression of *Ccr* NTD-YFP showed diffuse localization in *Eco* cells (Fig 2B), while the *Ccr* CTD in *Eco* cells showed robust rifampicin sensitive *Ccr* CTD-YFP foci (Fig 2B, Fig S10). These data show that the *Ccr* RNase E CTD is also sufficient for foci formation.

The conservation of the CTD across α-proteobacteria was then examined to explore how widespread BR-bodies are distributed. α-proteobacterial species contain a degenerate and poorly conserved CTD (Fig S6,S11). Despite the sequence heterogeneity, the CTD domains were predicted to contain a high degree of disorder with an alternating charge pattern across all α-proteobacterial (Fig S11). The disordered CTDs of different species were then assayed on their ability to promote BR-body assembly in *Ccr* when fused to the *Ccr* NTD. *Ccr* RNase E “replacement strains” where the RNase E-YFP hybrid fusions were the only expressed version of RNase E were generated. Here, control of the expressed form of RNase E can be performed using a xylose inducible copy of the wild type RNase E gene and a vanillate inducible copy of the mutant RNase E gene (Thanbichler et al., 2007) (Fig S12). Despite different sequence composition, the α-proteobacterial *Sme* or *Atu* CTDs robustly supported BR-body assembly when fused to the *Ccr* NTD (Fig 2C). The γ-proteobacterial *Eco* CTD which contains a membrane targeting signal could generate small weak foci that are localized at the cell membrane like the localization pattern observed in *Eco* cells (Strahl et al., 2015) (Fig 2C). These data show that robust cytoplasmic RNase E BR-bodies are a modular property of the α-proteobacterial CTDs, while those of γ-proteobacteria form small foci along the membrane as previously observed (Strahl et al., 2015).

Many eukaryotic proteins that assemble into LLPS droplets, such as those in p-bodies or stress granules, are also able to assemble LLPS droplets *in vitro* (Elbaum-Garfinkle et al., 2015; Lin et al., 2015). The intrinsically disordered regions of these proteins and often both necessary and sufficient for *in vitro* droplet formation and we similarly observed that the intrinsically disordered CTD of *Ccr* RNase E was able to form LLPS droplets *in vitro* (Fig 2D, S13). The CTD droplets were sensitive to ionic strength as observed with LLPS droplets of other membrane-less organelles (Lin et al., 2015) (Fig 2D, S13-14). Droplet assembly occurs at NaCl concentrations below 200 mM and requires protein concentrations of at least 12.4 μM (Fig 2D, S14). The strong dependence of the RNase E phase boundary on the salt concentration (Fig 2D) and patterns of alternating charged residues (Fig 2A,S9) suggests that electrostatics play an important role in the intermolecular RNase E interactions facilitating self-assembly into droplets. Because *in vivo* RNase E foci required the presence of RNA it was tested whether addition of RNA could induce LLPS of the CTD-YFP protein *in vitro*. Addition of 5-175 μg/mL poly-A RNA at a low concentration (12.4 μM) of CTD-YFP induced LLPS droplets and broadened the range in salt concentration over which they assemble (Fig 2D, S15-16). Interestingly, addition of excess poly-A RNA above 175 μg/mL prevented the assembly of droplets (Fig S15). This is likely due to stoichiometric interactions between the protein and RNA as observed in other systems (Lin et al., 2015). This observed inhibition of droplet formation at high RNA concentrations suggest that droplet formation may involve interactions between a RNA molecule and multiple copies of the RNase E protein. Taken together, these *in vitro* experiments indicate that CTD-YFP droplet formation likely involves both electrostatic protein-protein interactions and protein-RNA interactions as droplet formation is sensitive to both salt and RNA concentrations.

To investigate whether *in vivo* foci fusions display liquid-like properties, time-lapse microscopy was performed on the CTD-YFP protein in *Eco* cells. As highlighted, *Ccr* CTD-YFP foci can fuse together and quickly collapse into a spherical shape increasing the size of the YFP foci (Fig 2E). Rapid change of the foci shape is indicative of a liquid droplet while solid protein aggregates are rigid and resist deformation. Taken together, these experiments suggest that *Ccr* CTD-YFP have the hallmark characteristics of LLPS droplets both *in vitro* and *in vivo* and that their assembly is controlled by association with RNA. Therefore, we term these RNase E LLPS droplets <u>b</u>acterial <u>R</u>NP bodies (BR-bodies).

### RNase E recruits RNA degradosome components into BR-bodies

In addition to the alternating charge blocks, the CTD is also the primary scaffold for the RNA degradosome protein complex. In *Ccr*, the RNA degradosome complex contains RhlB, Aconitase, RNase D, and PNPase under logarithmic growth conditions (Fig 3) (Hardwick et al., 2011; Voss et al., 2014). To differentiate if the degradosome proteins localize in BR-bodies with RNase E or outside the bodies to inhibit RNase E assembly into bodies, two fluorescent fusions were generated for RhlB and Aconitase. Both RhlB-CFP and Aconitase-Chy fusions showed patchy foci within the cytoplasm with rifampicin sensitivity (Fig S17-18) suggesting RhlB and Aconitase likely localize in BR-bodies. To confirm that these RhlB-CFP and Aconitase-Chy foci were in BR-bodies colocalization with RNase E-YFP was assayed. RNase E-YFP foci were highly correlated with aconitase-Chy or RhlB-CFP foci (Fig 3), suggesting that these degradosome proteins are indeed assembled into the BR-bodies.

**Figure 3.**
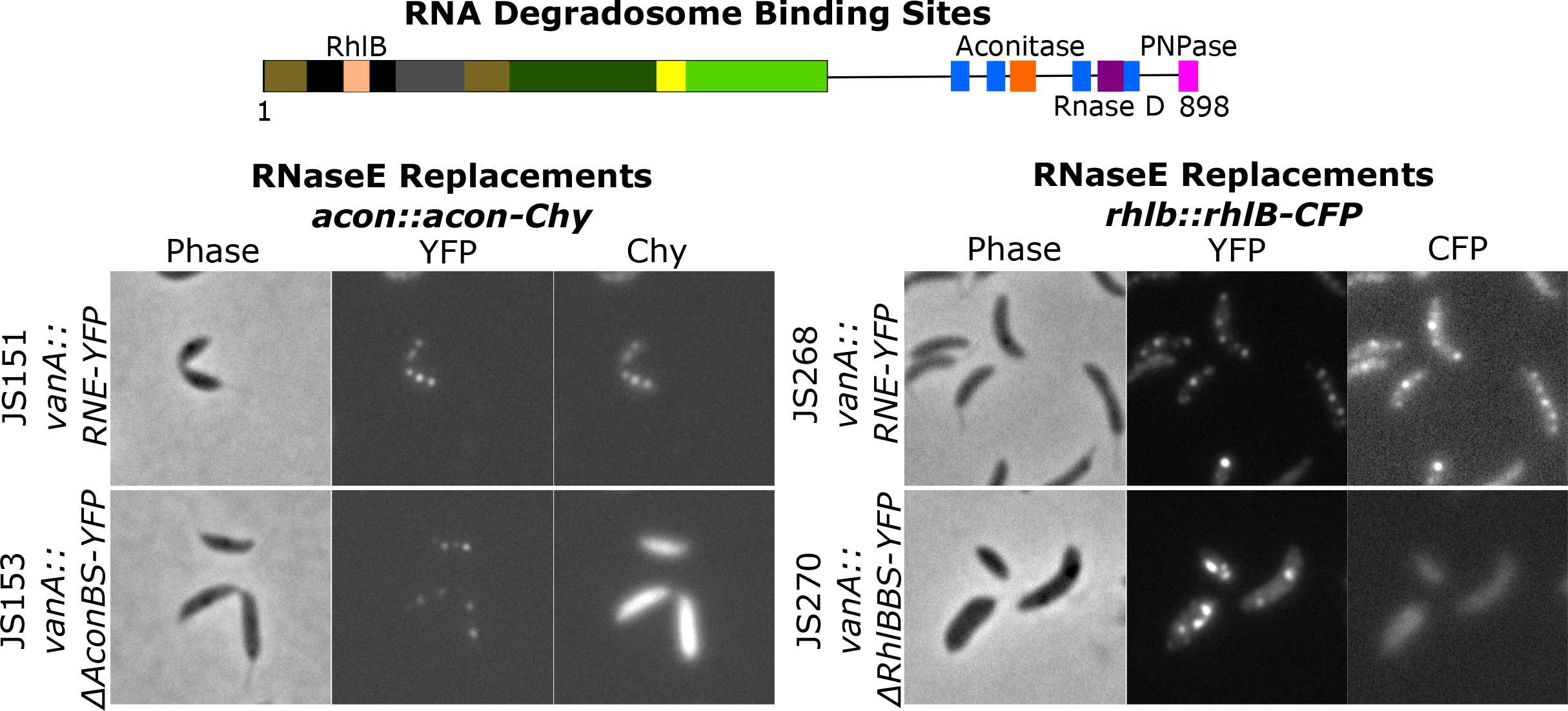
RNase E recruits degradosome components into BR-bodies. Degradosome binding site architecture for the *Ccr* RNase E protein with the RNA degradosome binding sites highlighted (Hardwick et al., 2011; Voss et al., 2014). Replacement strains harboring YFP tagged (RNE= full length, ΔAconBS=aconitase binding site deletion, ΔRhlBBS=RhlB binding site deletion) variants under control of the vanillate promoter and a natively tagged copy of acon-Chy (JS151 and JS153) or RhlB-CFP respectively (JS268 and JS270). All strains were grown under replacement conditions and imaged in log phase on a M2G 1.5% agarose pads.

We therefore hypothesized that BR-bodies may either assemble with RNase E forming a core scaffold of the BR-body that recruits the other degradosome proteins into the BR-body, or that RNase E together with the degradosome proteins form the core needed to assemble. To investigate the dependency of RNase E and the degradosome components on BR-body assembly, RNase E was depleted from the cells and subcellular localization of RhlB-CFP and Aconitase-Chy was examined. Localization of RhlB-CFP and Aconitase-Chy into cytoplasmic foci were both dependent on the presence of RNase E, as depletion of RNase E from cells led to a complete loss in RhlB or Aconitase foci (Fig S17-18). To test the dependency of the degradosome proteins on RNase E assembly into BR-bodies, RNase E deletion mutants in each of the respective degradosome protein binding sites were generated and their subcellular localization was assayed (Hardwick et al., 2011). In the aconitase and RhlB binding site deletion strains (ΔAconBS and ΔRhlBBS), the RNase E-YFP assembled robust BR-bodies, however, the aconitase-mChy and RhlB-CFP were diffuse across the cell body (Fig 3, Fig S19), suggesting that *in vivo* assembly of RNase E into BR-bodies does not depend on these degradosome proteins. In addition, deletion of the PNPase binding site (ΔPnpBS) or RNase D binding site (ΔRNDBS) also yielded RNase E proteins that maintained the ability to assemble BR-bodies (Fig S20-21). The ΔPnpBS and ΔAconBS mutants yielded slightly lower fluorescent protein levels (Fig S20), possibly due to lower protein stability or expression, but still maintained ability to form BR-bodies. Interestingly, cells expressing the ΔRhlBBS variant showed an increase in the number of BR-bodies per cell, however, these cells also contained a significant increase in cell size with apparent defects in cell division (Fig S20). As any single binding site mutant tested maintained the ability to assemble BR-bodies a ΔDBS (degradosome binding site) mutant (ΔAconBS/ΔRndBS/ΔPnpBS triple mutant) was generated that lacks all the degradosome binding sites in the CTD (Fig 20-21). The ΔDBS mutant yielded BR-bodies which were more diffuse but were similar in abundance to the wild type (Fig S20-21). In summary, deletion of any single degradosome binding site or a triple deletion mutant of all three degradosome binding sites in the CTD maintained the ability to assemble BR-bodies. This suggests that RNase E plays a major role in scaffolding BR-body assembly and allows recruitment of other degradosome components into BR-bodies.

### BR-bodies play a role in the *Ccr* stress response

In yeast, P-bodies and stress granules are increased rapidly upon cell stress (Brengues et al., 2005; Ramachandran et al., 2011; Shah et al., 2013). To test if BR-bodies are induced upon stress *Ccr* cells were subjected to several acute stresses and the subcellular distribution of RNase E-YFP was visualized. Like eukaryotes, several stresses led to an increase in the number of *Ccr* RNase E-YFP BR-bodies per cell and a decrease in diffuse RNase E-YFP signal including 10% ethanol, 30’ 5mM EDTA, and 5’ 42°C heat shock (Fig 4A, Fig S22). Washing away the ethanol with fresh M2G resulted in rapid disassembly of the BR-bodies, suggesting that the stress-induced BR-bodies dynamically disassemble after relieving the stress (Fig S23). Other stresses led to only minor increases in the number of *Ccr* RNase E-YFP BR-bodies including 5’ 0°C cold shock, 10mM H_2_O_2_, 110mM Sucrose, 30’ carbon starvation, stationary phase, and 200mM NaCl (Fig S22).

**Figure 4.**
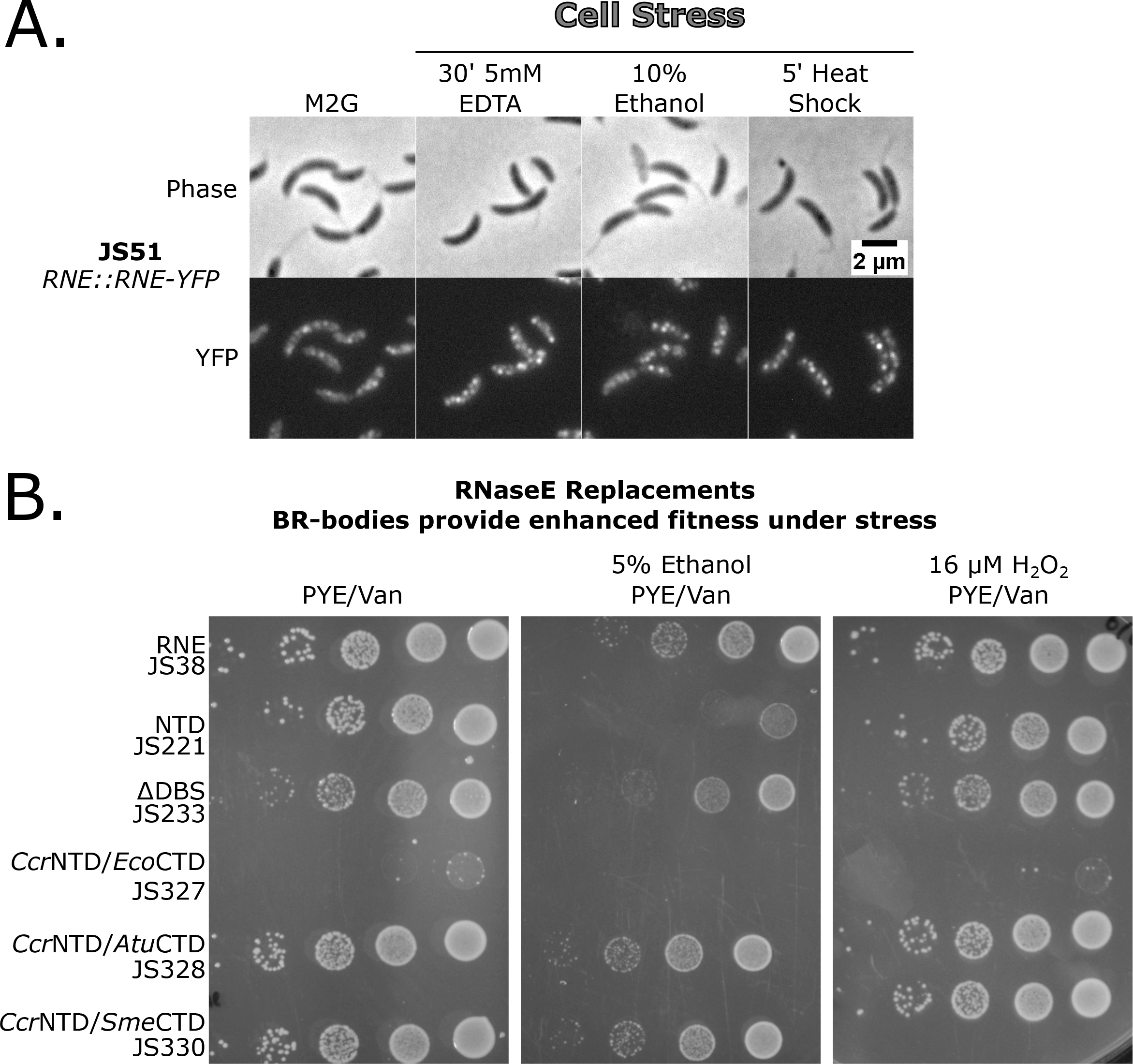
BR-body assembly is increased under specific stresses and provides increased stress tolerance. A.) Cells containing the RNase E-YFP fusion (JS51) were subjected to cell stresses and then imaged on M2G 1.5% agarose pads. The cells were treated with the stressor as indicated, gently mixed, spotted on the agarose pad, then immediately imaged. B.) BR-bodies provide enhanced survival under cell stress independent of degradosome formation. RNase E Replacement strains were grown under replacement conditions, diluted to the same OD600, and 1:10 serial dilutions were plated on PYE plates containing the selectable markers in the presence or absence of ethanol or H_2_O_2_ stresses.

The fitness of an RNase E replacement strain expressing either the full length *Ccr* RNase E or the NTD which lacks the ability to assemble BR-bodies were assayed. Because the CTD contains three degradosome protein binding sites, fitness of the ΔDBS replacement strain was also assayed. This allows the distinction of effects resulting from the lack of complete degradosome formation from those resulting from a lack of BR-body assembly. Under normal growth conditions, only subtle differences in growth were observed between the full length, NTD, and ΔDBS constructs (Fig 4B) suggesting that BR-body assembly and degradosome assembly are not required for log growth. Additionally, *Ccr*NTD/*Atu*CTD and *Ccr*NTD/*Sme*CTD assembled into cytoplasmic BR-bodies (Fig 2C, Fig S24) and grew normally, however, the *Ccr*NTD/*Eco*CTD strain which localizes RNase E to the inner membrane was extremely sick. This suggests that mislocalization of RNase E to the inner membrane does not support robust growth in *Ccr* cells.

As the BR-bodies are induced upon certain stress conditions (Fig 4A), growth of these strains under strong BR-body inducing conditions (ethanol stress) and weak BR-body inducing conditions (H_2_O_2_ stress) were tested. Under strongly inducing conditions the strain expressing the NTD showed marked defects in growth on plates containing 5% ethanol as compared to cells expressing the full-length version of RNase E (Fig 4B, Fig S25). The stress sensitivity was specific to conditions that increases BR-body assembly, as growth on plates containing 16 μM H_2_O_2_ only minimally lowered growth of cells expressing the NTD (Fig 4B). Interestingly, the defect in cell growth in ethanol stress was not due to the lack of degradosome formation, as the ΔDBS triple mutant lacking the ability to recruit multiple degradosome components was able to grow equivalent to wild type (Fig 4B). Additionally, *Ccr*NTD/*Atu*CTD and *Ccr*NTD/*Sme*CTD constructs, which are able to form cytoplasmic BR-bodies (Fig 2C, Fig S24), but assemble with different degradosome components (Zhang and Hong, 2009), were also resistant to ethanol stress. This suggests that BR-bodies give *Ccr* cells a fitness advantage under stress independent of the RNA degradosome proteins.

### RNase E BR-body disassembly requires RNA cleavage

Time-lapse microscopy was performed on live cells on M2G agarose pads to examine the dynamics of RNase E-YFP foci. Under normal log-growth conditions, *Ccr* RNase E-YFP assembles dynamic BR-bodies that assembled and disassembled within the 1-minute time scale at many locations along the long cell axis (Fig 5A). Upon treatment of cells with rifampicin to deplete cells of mRNA, RNase E-YFP BR-bodies were dramatically diminished (Fig 1A, Fig 5A). However, a small number of transient BR-bodies were observed that rapidly disassembled within one minute (Fig 5A). To test if the cleavage of mRNA was responsible for the rapid disassembly of BR-bodies, two separate active site mutants (ASMs) in the DNase I region of RNase E were generated. One mutant is a deletion of active site residue N362 (JS138) and the other has D360N/D403N mutations (JS299), in both cases disrupting residues that chelate the catalytic Mg^2+^ ion (Callaghan et al., 2005). Both ASM variants displayed an increased number of BR-bodies when compared to the wild type RNase E (Fig S26). The dynamics of the ASM variants were then monitored in an RNase E-YFP replacement strain where expression can be switched from the wild type RNase E to the catalytically inactive mutant using independent xylose and vanillate inducible promoters (Fig S12). The ASM-YFP replacement strains both assembled long lived foci that were stable across the entire 10-minute time lapse (Fig 5B, S27). A replacement strain of the CTD-YFP construct, which assembles robust BR-bodies, but lacks a catalytic domain also showed stable foci across the entire duration of the time-lapse and events of LLPS droplets fusing and growing were also observed (Fig 5B). These data suggest that mRNA cleavage is required for the dynamic disassembly of BR-bodies in *Ccr*.

**Figure 5.**
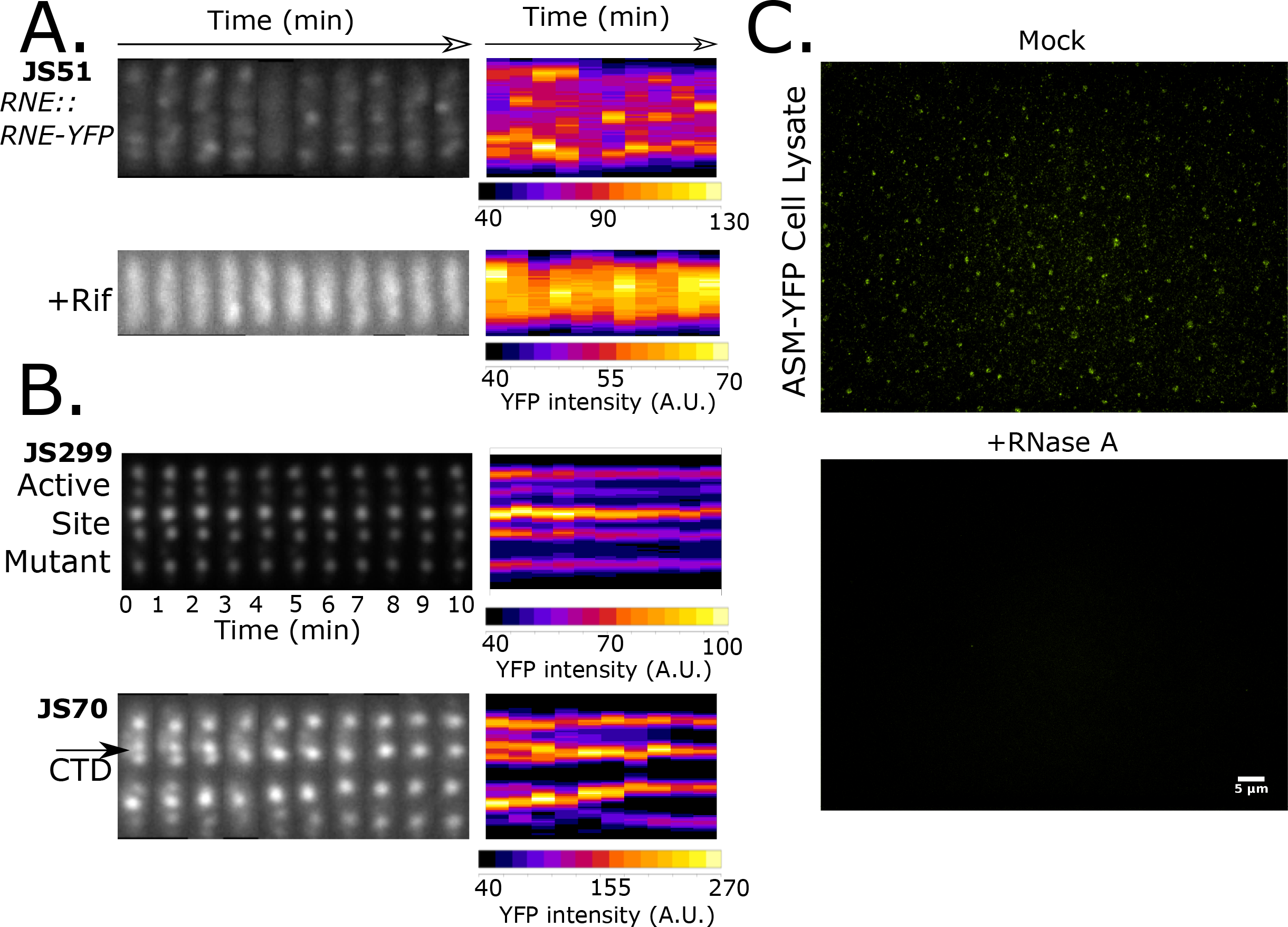
BR-bodies are dynamic and require RNA cleavage for disassembly. A.) *Ccr* RNase E-YFP (JS51) localization patterns were monitored by time-lapse microscopy of single cells grown on a M2G 1.5% agarose pad. Cells were either grown in M2G and spotted on the pad, or grown in M2G media and preincubated with rifampicin (100 μg/mL) for 30 min prior to spotting on the pad. All images were taken in 1 minute intervals. A representative time-lapse from a cell is shown for each condition with the straightened cell images placed side by side (kymographs generated via microbeJ). A heat map representation is plotted on the right of the same cell with its average YFP intensity across the long axis of the cell. B.) The active site mutant (JS299) is able to bind mRNA, but mutation of the residues involved in chelation of a catalytic Mg^2+^ ion renders it inactive (Callaghan et al., 2005) while the CTD variant (JS70) lacks the catalytic domain entirely. Time lapse was performed in the *Ccr* ASM and CTD replacement strain backgrounds under conditions where the ASM-RNase E or the CTD is the sole expressed form of RNase E. Black arrow indicates a BR-body fusion event. C.) Lysates of the ASM-YFP replacement strain (JS138) were prepared in a replacement strain background under conditions where the ASM-RNase E is the sole expressed form of RNase E. Lysates were then prepared and either treated with 0.1mg/mL RNase A or mock treated and imaged on a glass microscope slide.

As rifampicin treatments and *in vitro* LLPS droplet assays suggest that RNA is required to assemble BR-bodies and the cleavage activity of RNase E is required to disassemble them, the direct dependence of BR-body assembly on RNA was assayed in a cell lysate. Here a lysate of the replacement strain expressing only the ASM-YFP (JS138) was generated since it lacks the ability to cleave RNA substrates and assembles long lived foci that were predicted to remain stable after cell lysis. In ASM-YFP (JS138) lysates, many BR-bodies remained associated after cell lysis for the duration of our lysis and purification procedure (Fig 5C). After treating the lysate with RNase A for 30 minutes the BR-bodies were no longer detectable (Fig 5C) while mock treated lysates still contained BR-bodies, further supporting the findings that BR-body formation is regulated by RNase E-RNA interactions.

### Ribosomes and BR-bodies compete for mRNA substrates and affect decay

Eukaryotic mRNAs have been shown to go down competing paths, ribosomes can engage on the mRNA for translation or P-bodies can engage untranslated mRNA leading to degradation (Brengues et al., 2005). To determine if BR-bodies and ribosomes compete for mRNAs in *Ccr* the number of BR-bodies per cell was assayed under conditions known to alter the amount of translating and non-translating mRNA in the cell. When cells are depleted for mRNAs by rifampicin treatment, the number of BR-bodies per cell are reduced (Fig 1A, Fig 6A). Addition of translation elongation inhibitors chloramphenicol or tetracycline are known to accumulate ribosomes on mRNAs, leading to a decrease in the amount of untranslated mRNAs in the cell (Oh et al., 2011). Correspondingly, both chloramphenicol and tetracycline led to a reduction in the number of BR-bodies per cell (Fig 6A). Conversely, puromycin prematurely cleaves the peptide chain releasing the ribosome subunits and mRNA (Azzam and Algranati, 1973), therefore increasing the amount of untranslated mRNA in the cell. Consistent with the competition model, addition of puromycin led to an increased number of BR-bodies per cell (Fig 6A).

**Figure 6.**
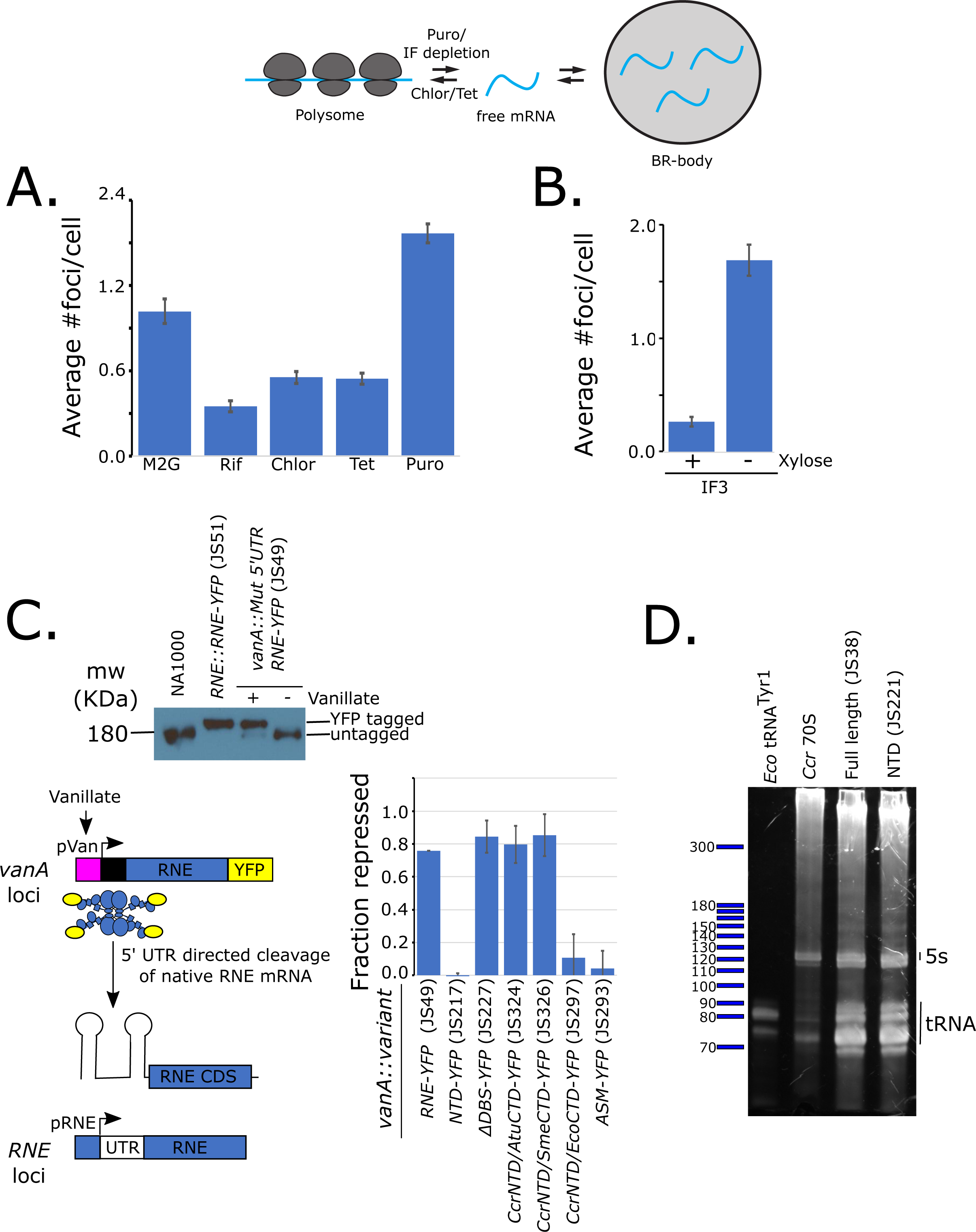
BR-bodies compete with ribosomes for untranslated mRNAs and affect mRNA turnover. A.) Number of JS51 RNase E-YFP foci per cell measured upon different acute antibiotic treatments measured using microbeJ. Concentrations used were 100μg/mL rifampicin for 30’, 200μg/mL chloramphenicol for 15’, 40μg/mL tetracycline for 15’, and 150μg/mL Puromycin for 30’. Error bars represent standard error. B.) Number of RNase E-YFP foci per cell measured upon depletion of translation initiation factor 3 measured using microbeJ. Gene encoding translation Initiation factors 3 under the sole control of the xylose promoter were grown in PYE/Kan/Xylose, then either maintained in xylose or grown in media lacking xylose for 8 hours to deplete the protein. Error bars represent standard error. C.) RNase E autoregulation assay where RNase E-YFP lacking the *Ccr* 5’ UTR expressed from the vanillate loci digests the 5’ UTR from the RNase E mRNA expressed from the RNE loci on the chromosome. Protein levels of the native RNase E were measured by western blot using an α-RNase E antibody. Protein levels of the untagged native RNase E in each strain tested was normalized to the protein level in NA1000 cells. D.) Total RNA from *Ccr* replacement strains was separated on an 8% acrylamide denaturing gel (1X TBE 7M Urea) and stained with sybr gold. An *in vitro* transcript of the mature *E. coli* tRNA^Tyr1^ (84 nt), purified 70S *Ccr* ribosomes, and 10bp ladder were loaded as controls.

The rate of cellular translation can also be controlled by altering the protein level of the cell’s translation initiation factors. Here, the gene for initiation factor IF3 was placed under the sole control of the xylose promoter, allowing depletion of IF3 upon switching the cells to media lacking xylose. The IF3 depletion strain showed increased abundance of BR-bodies per cell when switched to media lacking xylose for 8 hours (Fig 6B). The increase of BR-body number per cell concomitant with the decrease in the level of translation initiation further supports the model of competition. Growth under different nutrient conditions is also known to alter the cellular rates of mRNA translation. Polysome profiling showed that in *Ccr* cell growth in minimal medium (M2G) leads to slower rates of translation than growth in a rich growth medium (PYE) where translation rates are faster (Fig S28). Between these two conditions an inverse correlation in the number of BR-bodies per cell (Fig S28) was observed, again suggesting that *Ccr* BR-bodies may be more highly induced when translation rates are low. The observed inverse relationship between BR-body abundance and cellular translation rates across conditions suggests that BR-bodies and ribosomes compete for cellular mRNA substrates.

To examine the effects of BR-bodies on mRNA decay activity of RNase E, the ability of RNase E to cleave its 5’ UTR (Jain and Belasco, 1995) was assayed. In *Ccr* cells a YFP tagged variant of RNase E expressed from the *vanA* locus with an exogenous 5’ UTR digests the RNase E mRNA through its native 5’ UTR, lowering the steady state level of the native untagged protein (Fig 6C). In this assay, expression of the full length YFP tagged *Ccr* RNase E led to 76% (± 9.8% S.E.M. n=4) repression of the native protein level, while a control strain expressing the catalytically null active site mutation led to undetectable levels of repression (4.2% ± 8.9% S.E.M. n=3). Interestingly, expression of the NTD mutant which cannot form BR-bodies led to a complete loss in repression (0% ± 11% S.E.M. n=6), while the ΔDBS strain which assembles BR-bodies but lacks the ability to recruit a complete degradosome showed a robust 84% repression (± 13% S.E.M. n=2). The hybrid variants containing either CTDs from *Atu* or *Sme* both led to robust repression 80% (± 14% S.E.M. n=2) and 85% (± 11% S.E.M. n=2) respectively, while the *Eco* hybrid which associates with the inner membrane led to very low levels of repression (11% ± 6.0% S.E.M. n=2). These results show that the ability to assemble a cytoplasmic BR-body is important for *Ccr* RNase E mRNA degradation activity independent of RNase E’s ability to recruit the degradosome components into the body. Interestingly, mature 5s rRNA and tRNAs accumulated normally in the variant lacking the CTD (Fig 6E), suggesting that stable rRNA and tRNA processing does not require BR-body formation.

## Discussion

### Role of liquid-liquid phase separation in bacterial cell organization

Bacteria were once thought of as amorphous “bags of enzymes”, due to the general lack of membrane bound organelles and discrete subcellular niches. It has become increasingly apparent that the bacterial cytoplasm uses many mechanisms for organizing cellular processes including cytoskeletal proteins for directing cell wall synthesis (Garner et al., 2011), polarly localized scaffolding complexes for regulating chromosome segregation (Holmes et al., 2016), or protein shelled microcompartments like carboxysomes that house CO_2_ fixation (Cameron et al., 2013). Studies of mechanisms that organize the bacterial cytoplasm have revealed general mechanisms of diffusion and capture, cell wall curvature, nucleoid occlusion, protein shell assembly, and affinity for membrane and/or cell-wall components for establishing subcellular zones to organize cellular biochemistry (Barry et al., 2014; Laloux and Jacobs-Wagner, 2014; Rudner and Losick, 2010). This report extends these bacterial organization mechanisms to include liquid-liquid phase separation of membrane-less “droplet organelles” with capacity to partition mRNA degrading enzymes and mRNA substrates into BR-bodies. The intrinsically disordered CTD of *Ccr* RNase E is necessary and sufficient for LLPS droplet formation *in vivo* and *in vitro* (Fig 2) and contains patches of positively and negatively charged residues similar to eukaryotic LLPS proteins (Nott et al., 2015). These charged patches allow the BR-bodies to self-assemble into droplets through electrostatic interactions and the positively charged patches may also facilitate binding to RNA, thereby allowing RNase E to increase its local concentration to accelerate the rate of self-assembly (Fig 7). The dynamic assembly and disassembly of BR-bodies (Fig 5) is different than other microcompartments such as carboxysomes, which form static protein shells, and this dynamic control of BR-body assembly may be utilized to regulate enzymatic activity of the proteins inside. Interestingly, 129 other proteins with “patchy foci” subcellular localization patterns have been observed in *Ccr* (Werner et al., 2009), including aconitase which functions in the TCA cycle (Fig 3), suggesting that the LLPS mechanism of organization may be more broadly utilized across other biochemical pathways in this bacterium. Future studies are needed to understand the breadth of how and when membrane-less droplet organelles are assembled, regulated, and used in bacteria, including the characterization of polymer-like properties of other intrinsically disordered self-assembled signaling hubs such as the PopZ scaffolding complex in *Ccr* (Holmes et al., 2016).

**Figure 7.**
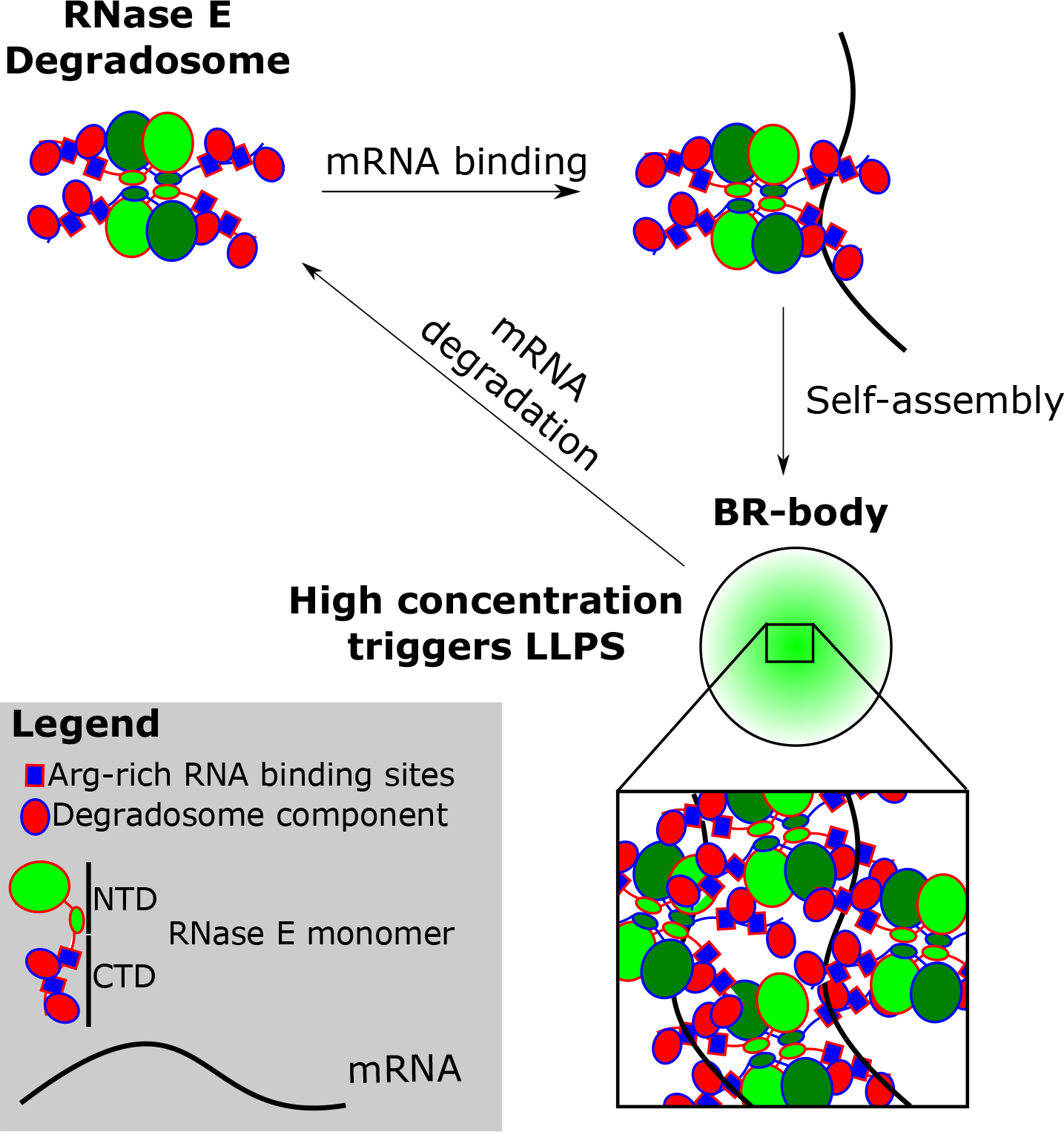
Model of *Ccr* RNase E BR-body assembly and disassembly. Free RNase E tetramers scaffold the RNA degradosome protein complex. In the presence of untranslated mRNA, RNA degradosomes can bind to the mRNA and when the concentration of bound RNA degradosomes is high, liquid-liquid phase separated (LLPS) droplets self-assemble via the alternating charged-blocks in the intrinsically disordered CTD of RNase E. Upon initial RNA cleavage by RNase E, degradosome associated factors rapidly facilitate mRNA degradation allowing the RNase E degradosome complexes to be disassembled from the BR-body.

### Diverse subcellular architectures of bacterial mRNA turnover

Bacterial RNase E have two distinct modes of subcellular organization across species; a membrane tethered organization in γ-proteobacteria and a cytoplasmic organization in α-proteobacteria that assembles robust BR-bodies described here. The intrinsically disordered CTD acts as a modular unit controlling the subcellular localization pattern (Fig 2). The membrane localized mode of subcellular localization in γ-proteobacteria facilitates rapid turnover of mRNAs encoding inner-membrane proteins (Moffitt et al., 2016) and here the cytoplasmic localization of α-proteobacteria appears to allow BR-bodies to assemble. In *Eco* the membrane tethering sequence and mRNA are required to assemble small RNase E foci (Strahl et al., 2015), yet it is not known whether they assemble into LLPS droplets and what limits the *Eco* foci size. Interestingly, the hybrid *Ccr*NTD/*Eco*CTD which localizes to the membrane cannot support growth of *Ccr* cells, reducing growth even compared to a strain lacking the CTD entirely (Fig 4B), suggesting α-proteobacterial bacterial cells are strongly adapted to using a cytoplasmic localization strategy. Interestingly, under anaerobic growth, Enolase, a glycolytic component of the *Eco* degradosome, facilitates *Eco* RNase E to switch to a diffuse cytoplasmic localization pattern (Murashko and Lin-Chao, 2017) that is needed to switch *Eco* cells to a program of filamentous growth. While these findings are indicative that RNase E substrate specificity and activity can be regulated by its subcellular distribution, the biochemical consequences of each strategy remain poorly understood.

### BR-bodies – multifunctional RNP bodies

In this study α-proteobacterial cells were shown to contain LLPS BR-bodies, providing evidence that some bacterial species and eukaryotes use a similar method to organize their cellular mRNA turnover machinery (Fig 7). Eukaryotic P-bodies or stress granules and BR-bodies require RNA to assemble, require proteins with intrinsically disordered regions, are induced upon cell stress, and contain the cell’s major mRNA degrading nuclease and other mRNA decay factors. Interestingly, super resolution imaging of *Ccr* BR-bodies by RNase E-YFP found their size (between 0.1 to 0.4 μM) (Bayas et al., 2018) are on the same order of magnitude as those of p-bodies or stress granules (Wheeler et al., 2016). Additionally, 3D single molecule particle tracking of *Ccr* RNase E-YFP found that diffusion of RNase E-YFP molecules were confined, most likely within LLPS droplets, and the number of confined RNase E-YFP molecules dropped after rifampicin treatment (Bayas et al., 2018). BR-bodies and P-bodies have also been shown to compete against ribosomes for mRNA substrates (Fig 6) (Brengues et al., 2005) suggesting BR-bodies likely accumulate non-translating mRNAs. BR-bodies appear to play an important role in mRNA turnover in *Ccr* cells as mutants lacking the ability to form BR-bodies were found to be deficient in decay of the RNase E mRNA independent of RNA degradosome formation (Fig 6). How BR-bodies facilitate mRNA decay is still a mystery, but it remains possible that BR-body assembly helps concentrate RNase E and mRNAs increasing the reaction rates of the decay process. Non-stop decay factors such as tmRNA and SmpB were also shown to localize into discrete foci in *Ccr*, suggesting that non-stop decay and ribosome rescue may be performed in BR-bodies (Russell and Keiler, 2009). Yet many established functions of eukaryotic P-bodies or stress granules are still unclear in bacteria. Does bacterial mRNA 5’ PPP to 5’ P “decapping” by RppH which can stimulate RNase E activity occur in BR-bodies? Can BR-bodies function to store mRNAs or are they used mainly for mRNA turnover? Are bacterial small RNA regulatory machinery accumulated in BR-bodies like RISC complexes and miRNAs are in P-bodies?

Like P-bodies and stress granules, BR-bodies accumulate rapidly upon certain stresses (Grousl et al., 2009; Shah et al., 2013; Teixeira et al., 2005). More similar to P-bodies, BR-bodies appear to rapidly disassemble when the stress is removed (Fig S23), whereas stress granules have slower assembly/disassembly dynamics. Replacement of RNase E with a variant lacking the CTD (JS221) that is unable to assemble BR-bodies and led to a significant fitness defect to *Ccr* cells when grown under strongly inducing stress (ethanol stress) and only a mild fitness defect under weakly inducing stress (H_2_O_2_ stress) (Fig 3B). Importantly, increased cell fitness under stress was independent of the degradosome formation defects in the CTD deletion, as the ΔDBS mutant (JS233), which can assemble BR-bodies while lacking all degradosome binding sites in the CTD, grew similarly to wild type under ethanol and H_2_O_2_ stresses. This data suggests that the enhanced stress resistance is provided by BR-body assembly itself which may play important roles in related pathogenic α-proteobacteria, such as *Brucella abortus*, to survive in the presence of the immune system.

While the eukaryotic nuclease Xrn1 has its main function in cytoplasmic mRNA decay, *Ccr* RNase E functions in mRNA decay in addition to rRNA processing (Hardwick et al., 2011) and tRNA processing (Ow and Kushner, 2002) all within the nucleoid filled cytoplasm. BR-bodies are a multifunctional “swiss army knife,” RNP body, encompassing functions of P-bodies, stress granules, and the nucleolus within a single entity. In another study, BR-bodies were found to assemble at significant levels adjacent to both rRNA loci (Bayas et al., 2018) on the chromosome where RNase E may facilitate cotranscriptional 9s rRNA processing (Hardwick et al., 2011) bearing similarity to the LLPS eukaryotic nucleolus (Courchaine et al., 2016). Additionally, RNase D is a component of the *Ccr* degradosome and it plays a role in tRNA 3’ end maturation, suggesting that tRNA processing may occur in BR-bodies as well. Surprisingly, replacement of RNase E with the CTD deletion variant which cannot form BR-bodies still led to normal accumulation of mature 5s rRNA and tRNAs, suggesting that BR-bodies are not essential for stable RNA processing (Fig 4B, 6E).

BR-bodies can assemble with RNase E as the sole core protein component, which is simpler than that multi-component core of eukaryotic P-bodies where assembly *in vivo can occur* even in the absence of Xrn1 (Sheth and Parker, 2003). While a set of four degradosome proteins can purify with RNase E in stoichiometric amounts under normal growth conditions (Hardwick et al., 2011; Voss et al., 2014), when *Ccr* cells are grown in the cold, RhlB was shown to be displaced from RNase E and another DEAD box helicase RhlE and the transcription termination factor Rho were found associated (Aguirre et al., 2017). The changes in RNase E association across different environmental conditions suggesting that the composition of the BR-body may be regulated to control the cell’s changing needs of RNA processing/decay.

## Materials and Methods

### Cell growth

All *Caulobacter crescentus* strains used in this study were derived from the wildtype strain NA1000, and were grown at 28°C in peptone-yeast extract (PYE) medium or M2 minimal medium supplemented with 0.2% D-glucose (M2G) (Schrader and Shapiro, 2015). When appropriate, the indicated concentration of vanillate (5μM), Xylose (0.2%), gentamycin (0.5μg/ml), kanamycin (5μg/ml), chloramphenicol (2μg/ml), spectinomycin (25μg/ml), and/or streptomycin (5μg/ml) was added. Strains were analyzed at mid-exponential phase of growth (OD 0.3-0.6). Optical density was measured at 600 nm in a cuvette using a NanoDrop 2000C spectrophotometer. For acute drug treatment to deplete mRNA, the log phase cells were treated with Rifampicin (100μg/ml) for 30 minutes. Cell stress treatments were performed by incubating cells with the desired stress then immediately spotted on a 1.5% agarose pad and imaged or by incubating with the stress for the indicated time then spotting on the pad and imaging. Replacements strains containing a xylose inducible copy of RNase E and a Vanillate inducible test construct were first grown in media containing xylose overnight, then washed 3 times with 1mL growth media, and resuspended in growth media containing vanillate, diluted, and grown overnight (Fig S13). Log-phase cultures were then spotted on M2G 1.5% agarose pads for imaging. For serial dilution assays, cells were grown overnight in xylose containing media, washed 3 times, and diluted to OD 0.05. Serial dilutions were then performed and spotted on plates with indicated composition and incubated at 28°C.

*E. coli* strains were grown at 37°C and cultured in Luria-Bertani medium (L3522, Sigma), supplemented with the indicated concentration of kanamycin (30μg/ml) or Ampicillin (50μg/ml). For induction, BL21 DE3 cells were induced with isopropyl-3-Dthiogalactopyranoside (IPTG) (1μM) for 2 hours and TOP10 cells were induced with 0.0004% arabinose for one hour. Strains were analyzed at mid-exponential phase of growth (OD_600_ 0.3-0.6). Optical density was measured at 600 nm in a cuvette using a NanoDrop 2000C spectrophotometer.

*Agrobacterium tumefaciens* C58 cells were grown in Luria-Bertani medium. When appropriate, the indicated concentration of gentamycin (100μg/ml) was added. *Sinorhizobium meliloti* 1021 cells were grown in Luria-Bertani medium. When appropriate, the concentration of gentamycin (50μg/ml) was added. Strains were analyzed at mid-exponential phase of growth (OD 0.3-0.6). Optical density was measured at 600 nm in a cuvette using a nanodrop 2000C spectrophotometer.

### Cell imaging and analysis

For imaging, cells were immobilized on 1.5% agarose pads made with M2G medium on microscope slides (3051, Thermo Scientific) and images were collected using a Nikon Eclipse NI-E with CoolSNAP MYO-CCD camera and 100× Oil CFI Plan Fluor (Nikon) objective, driven by Nikon elements software. The filter sets used for YFP, CFP and mCherry imaging were chroma 96363, 96361, and 96322 models respectively. For time lapse experiments, images were taken at 1-minute intervals with autofocus on over a maximum time of 10 minutes.

Cell image analysis was performed using microbeJ (Ducret et al., 2016). For automated foci detection we used the maxima foci function of microbeJ we manually adjusted the tolerance and Z-score settings to identify and outline foci on a test image then removed aberrant foci with area <0.01 μm^2^ and length >1 μM. The segmentation option was also used to split adjoined foci. For a given set of images we applied the same tolerance/Z-score parameters to quantify the number of foci/cell with a minimum of 50 cells.

### RNA extraction and analysis

For RNA extraction log-phase cells were rapidly pelleted at 14k rpm in a microcentrifuge and resuspending in 65°C Trizol as in (Zhou et al., 2015). RNA were run on 1× TBE/7M Urea denaturing PAGE gels and visualized with Sybr-Gold stain.

### Western blot

For the western blot, Log-phase cells were induced with vanillate (0.5mM) for 6 hours or left uninduced. Upon induction, the cells were pelleted and resuspended in 250μl of 1× laemmli buffer for each 1.0 OD_600_ unit. The samples were boiled at 95°C for 5 minutes then vortexed. 15μl of samples were loaded in a 10% SDS-PAGE gel and run at 200V for 90 minutes. Semi-dry transfer was done at 40 milli-amps for 1 hour after which the membrane was dried for 1 hour at room temperature (in the dark). The membrane was rewet in 100% methanol for 15 seconds and rinsed in Milli-Q water for 2 minutes. Blocking was done for 1 hour using 15ml of blocking buffer (1×PBS, 0.1% tween20, 0.05g/ml non-fat dry milk) at room temperature with gentle shaking. For primary antibody blotting, the membrane was submerged in (1:1000) dilution of the α-RNE antibody (Ben Luisi, U. Cambridge) in the same blocking buffer and shacked gently for 1 hour at room temperature. After washing the membrane 3 times with the same blocking buffer for 10 minutes the membrane was probed with secondary antibody (1:5000) anti-rabbit IgG secondary antibody (thermo-scientific) in blocking buffer. Secondary antibody incubation was done for 1 hour with gentle shaking at room temperature. The membrane was then washed 3 times, 10 minutes each, with 1× PBS buffer with gentle shaking. Following the wash, the membrane was placed in Pierce chemiluminescence substrates for 5 minutes and imaged on film.

### BR-body RNase digestion assay

Js138 (ASM) cells were grown in PYE medium supplemented with gentamycin (0.5μg/ml), kanamycin (5μg/ml), and 0.2% xylose for 15 hours. The next day, cells were washed three times with 1 ml of PYE and incubated in three dilutions in PYE supplemented with gentamycin (0.5μg/ml), kanamycin (5μg/ml), and vanillate (5μM and grown overnight (15 hours). The next morning, log-phase cells (OD 0.2-0.3) were harvested and pelleted at 16K×g at 4°C and then resuspended in 2ml of lysis buffer (35mM NaCl, 20mM Tris-HCl pH 7.04, 1mM EDTA-free protease inhibitor tablet, 1mM BME (freshly added), small pinch of Lysozyme and 0.8 U of RNase free DNase I) and flash frozen by dripping into liquid nitrogen. Cells were lysed in a mixer mill MM400 in 50mL chambers for 6 rounds at 15Hz for 2 minutes under liquid nitrogen. The lysate was thawed at room temperature and transferred into eppendorf tubes. The tubes were spun at 2000×g for 5 minutes at 4°C and the supernatant and pellet fractions were separated. 8 uls of the supernatant were either treated with 2μg of RNase A (2μl of 1mg/ml) or mock treated with 2μl of milliQ water and incubated at room temperature for 60 minutes. The entire 10μl reaction was then spotted on a glass microscope slide and imaged under a coverslip.

### CTD-YFP protein purification protocol

The C-terminal half of RNase E from *C. crescentus* (451-898) harboring an N-terminal His_6_ tag and a C-terminal YFP fusion was expressed from plasmid [pET28-RNE(CTD)-YFPC] that was transformed into chemically competent BL21 (DE3) cells before being plated onto selective LB-Miller media (50 μg/mL kanamycin plates) and incubated overnight at 37 °C. From a single colony, an overnight 60 mL LB-Miller culture (50 μg/mL kanamycin) was inoculated and incubated overnight at 37 °C. From this saturated culture, 6 L of LB-Miller media (50 μg/mL kanamycin) was inoculated and grown to mid-log phase (^~^0.5 OD_600_). Expression of RNase E (451-898) was induced with 1 mM isopropyl-β-D-thiogalactopyranoside (IPTG) for 4 hours at 37 °C. The cells were then collected at 4 °C, 4000 × *g*, for 30 minutes. The resulting pellet was washed with 50 mL 50 mM HEPES pH 8.0, 500 mM NaCl before being pelleted again at 4 °C, 4000 × *g*, for 30 minutes and stored at −80 °C.

Cells were thawed on ice and resuspended in 100 mL of lysis buffer (1 M NaCl, 20 mM Tris-Cl pH 7.4, 20 mM imidazole pH 7.0, 1 mM β-mercaptoethanol [βME], 20 U DNase I, and 0.1% Triton X-100, supplemented with SIGMAFAST™ protease inhibitor tablets [Sigma]). The cells were lysed by continuous passage through an Avestin EmulsiFlex-C3 at 15,000 psi for 30 minutes. Cell debris was pelleted by ultracentrifugation (4 °C, 20000 × *g*, 30 minutes). The supernatant was then incubated with 6 mL of a 50% slurry of Ni-NTA agarose at 4 °C for 2 hours. The Ni-NTA agarose was separated by a gravity column and washed with 30 mL wash buffer (150 mM NaCl, 20 mM Tris-Cl pH 7.4, 40 mM imidazole pH 7.0, 1 mM βME). RNase E (451-898) was eluted from the Ni-NTA agarose with 10 mL portions of elution buffer (150 mM NaCl, 20 mM Tris-Cl pH 7.4, 200 mM imidazole pH 7.0, 1 mM βME). The eluate was concentrated by a 30 kDa MWCO Amicon Centrifugal Filter Unit to 1 mL before being loaded onto a HiLoad Superdex 16/600 200 pg gel-filtration column (GE Healthcare) and eluted with storage buffer (150 mM NaCl, 20 mM Tris-Cl pH 7.4). Fractions containing RNase E were collected and concentrated using Amicon Centrifugal Filter Units (30 kDa MWCO) to ~30 mg/mL (M.W. = 80.77 kDa, ε_280_ = 55,600 M^-1^cm^-1^) before being aliquoted, flash-frozen in LN_2_, and stored at −80 °C.

### Liquid-liquid phase separated droplet assembly assay

RNase E CTD (451-898) protein aliquots were thawed on ice along with solutions of poly(A) and DNase/RNase-free water. All solutions were passaged through a 0.2 μm cellulose acetate syringe filter (VWR) and were stored in polypropylene 1.7 mL or 2.0 mL tubes (VWR). Prior to imaging, VistaVision microscope slides (VWR) were cleaned by rinsing in DNase/RNase-free water at ^~^70 °C before being soaked in RNase AWAY™ (ThermoFisher) overnight. The microscope slides were then rinsed three times in DNase/RNase-free water before being stored in 95% ethanol.

Solutions of RNase E (51-898), RNA, and NaCl were combined, diluted with sterile-filtered DNase/RNase-free water to various concentrations, and allowed to incubate on ice for no more than 1 hour prior to imaging. Microscope slides were blow-dried with air before 10 μL were pipetted onto the slides and covered with coverslips (VWR). All images were taken with an Eclipse Ti-E inverted microscope (Nikon) in both phase-contrast and fluorescent channels using a Plan Apo 100× objective. Phase diagrams were generated by yes/no scoring for droplet assembly based on fluorescent imaging.

## Author Contributions

JMS conceived the project. NA and JMS performed the cell biology experiments and generated the strains/plasmids. DTT and WSC performed the purification of CTD-YFP protein and *in vitro* LLPS droplet assays. JMS and NA wrote the paper.

## Acknowledgements

Research reported in this publication was supported by NIGMS of the National Institutes of Health under award number R35GM124733. The authors thank Wayne State University startup funds to JMS and University of Pittsburgh for startup funds to WSC. The authors thank O.C. Uhlenbeck for the T7 RNA polymerase plasmid, G.R. Bowman for the msfGFP plasmid, and B.F. Luisi for the α-RNase E antibody. The authors thank members of the Alcedo, Ansari, Meller, Beningo, and Higgs labs for sharing equipment and reagents. The authors thank O.C. Uhlenbeck, members of the Higgs lab, and C. Bayas for thoughtful discussion.

